# The human superior colliculus motor region does not respond to finger movements

**DOI:** 10.1101/2023.03.30.534871

**Authors:** Nikhil G Prabhu, Nicole Knodel, Marc Himmelbach

## Abstract

Electrophysiological studies in macaques and functional neuroimaging in humans revealed a motor region in the superior colliculus (SC) for upper limb reaching movements. Connectivity studies in macaques reported direct connections between this SC motor region and cortical premotor arm, hand, and finger regions. These findings motivated us to investigate if the human SC is also involved in sequential finger tapping movements. We analysed fMRI task data of 130 participants executing simple finger tapping from the Human Connectome Project (HCP). While we found strong signals in the SC for visual cues, we found no signals related to simple finger tapping. Assuming a differential effect owing to simple and complex finger tapping sequences, we searched for responses in the SC corresponding to complex finger tapping above simple finger tapping sequences. We observed expected signal differences in cortical motor and premotor regions, but our analyses provided no evidence for signals related to simple or complex finger tapping in the SC. Despite evidence for direct anatomical connections of the SC motor region and cortical premotor hand and finger areas in macaques, our results suggest that the SC is not involved in simple or complex finger tapping in humans.

## Introduction

Several electrophysiological studies in macaques demonstrated the involvement of the SC in visually guided arm movements (Werner, 1993; Werner et al., 1997a; Werner et al., 1997b; Stuphorn et al., 1999; Stuphorn et al., 2000). Philipp and Hoffmann (2014) reported the initiation and execution of arm movements upon electrical microstimulation of the SC in macaques. These observations were confirmed in humans by two fMRI studies on visually guided reaching (Linzenbold and Himmelbach, 2012; Himmelbach et al., 2013). A recent study found equivalent signals at this SC motor region for reaching to visual and tactile targets (Prabhu and Himmelbach, 2022). Cooper and McPeek (2021) reviewed the evidence for the involvement of the SC in body and limb movements across species from lampreys to humans. Their overview showed that a functional role of the SC in upper limb movements is not unusual from a phylogenetic perspective, with considerable variability across species.

Anatomical studies in macaques reported connections between the homologues of the human ventral and dorsal premotor cortices (PMv, PMd) and the upper limb motor regions of the SC (Fries, 1985; Borra et al., 2014; Distler and Hoffmann, 2015). In humans, PMv and PMd are involved in the execution of simple and complex finger tapping movements (Jäncke, Himmelbach et al., 2000; Jäncke, Loose et al., 2000; Horenstein et al., 2009; Ruspantini et al., 2011; Genon et al., 2017; see Witt et al., 2008 for a meta-analysis). We presumed that the SC also plays a role in finger tapping movements. This presumption was supported by findings in our earlier study on reaching movements where we observed signal increases in the SC limb motor region for a control condition that included button presses but no reaching movements (Prabhu and Himmelbach, 2022). In the current study, we used a publicly available dataset and a dedicated fMRI experiment to examine whether the previously identified SC upper limb motor region also reveals signals during finger tapping sequences. In the first part of our study, we used the task fMRI dataset from the Human Connectome Project (HCP, Elam et al. 2021). Since we did not find any finger tapping activation in the SC with the HCP dataset we conducted an fMRI experiment that analysed simple and complex finger tapping movements paced by visual or auditory cues.

### Human Connectome Project data

The HCP motor task had been modified from a study by Buckner et al. (2011). It consisted of right and left toe movements, right and left finger movements, and tongue movements. From the HCP dataset, we analysed 130 subjects looking for signals during finger tapping at the SC location where we observed a signal peak for a button press control condition in our earlier study (Prabhu and Himmelbach, 2022).

### Simple and complex finger tapping experiment

We found no evidence for SC involvement with simple finger tapping in the HCP data. Early studies on the premotor cortex reported differential signals for simple and complex finger tapping sequences (for an overview and meta-analysis please see Witt et al. 2008). Thus, we presumed that the null-finding in HCP data might be due to the use of simple finger tapping only and continued our search with an experiment including simple and complex finger tapping upon visual and auditory pacing cues. We measured and analysed 4 participants with each participant repeating the experiment four times on four different days with an exceptionally large number of blocks in multiple single-case replications. We reasoned that highly trained and experienced participants with a large number of trials would provide us with more reliable within-participant estimates of true effect sizes. Similar small-N approaches have been used earlier in fMRI (O’Connor et al., 2002; Wunderlich et al., 2005; Konen and Kastner, 2008; Meier et al., 2008; Konen et al., 2013; Spitschan et al., 2017; Wu et al., 2022; Fracasso et al. 2022). Please see Smith and Little (2018), Baker et al., (2021), and Ince et al., (2022) for recent discussions of and perspectives on small-N approaches.

## Results

### Human Connectome Project data

We first present the results of a second-level GLM analysis of right and left hand finger tapping from 130 subjects (*p <* 0.05, FWE-corrected). The task consisted of 10 finger movements over 12 s, with 2 blocks of left hand and 2 blocks of right hand finger tapping per run and 2 runs per subject. Finger tapping conditions are expected to show robust activation clusters in the primary motor cortex (M1), dorsal pre-motor cortex (PMd), and ventral pre-motor cortex (PMv) (please see Witt et al., 2008 for a meta-analysis). The crosshairs in figure 1 are centred on the global maxima for left and right M1, PMd, and PMv corresponding to right and left hand finger tapping respectively. These cortical results confirmed the validity of our approach and analysis with respect to the analyses and results that follow for the SC motor region.

**Figure 1:**
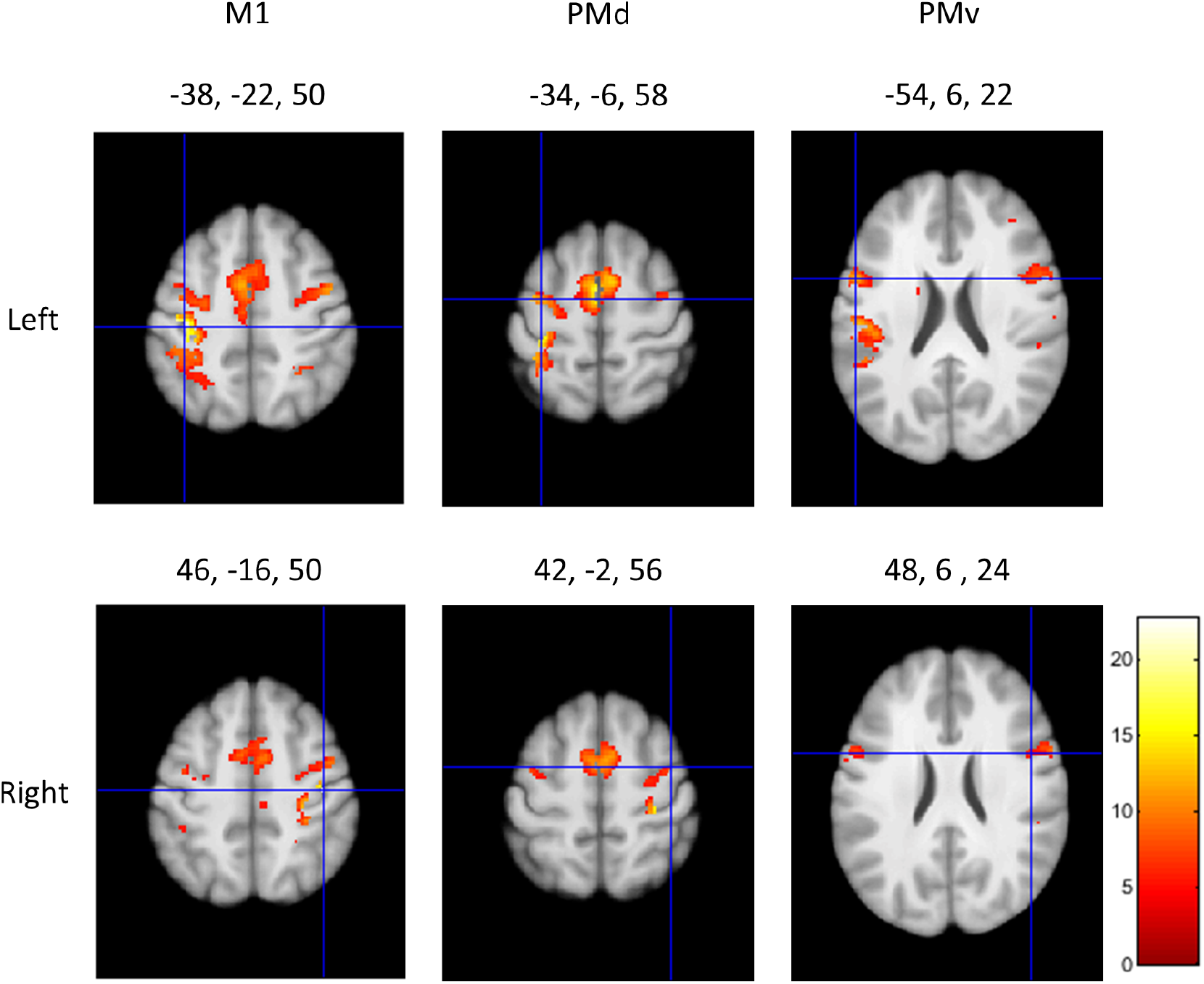
Cortical results for finger tapping from the HCP motor task dataset. Second-level GLM results centred on the peak coordinates in left M1, PMv, and PMd corresponding to right hand finger tapping (upper row), and right M1, PMv, and PMd corresponding to left hand finger tapping (lower row). The results have been thresholded at *p <* 0.05 (FWE-corrected). The colourbar shows the range of t-values.

We analysed GLM contrast estimates for visual cues, right hand, and left hand finger tapping at the peak location of button press activity that we observed in an earlier study (Prabhu and Himmelbach, 2022), i.e. at MNI coordinates -6, -28, -6 (volume of interest, VOI). We also included the corresponding location in the right SC at 6, -28, -6 as the HCP dataset includes right and left hand finger tapping. We found a large, positive signal for the visual cues but no positive signal for right and left hand finger tapping. For the motor conditions the 90% confidence intervals clearly overlapped with 0 at both the locations (Figure 2). The figure shows 90% confidence intervals for a two-sided test. Please note that our hypothesis was uni-directional, hence, we considered the results to be significant if the lower bound of the interval which amounted to 95% confidence, was above zero.

**Figure 2:**
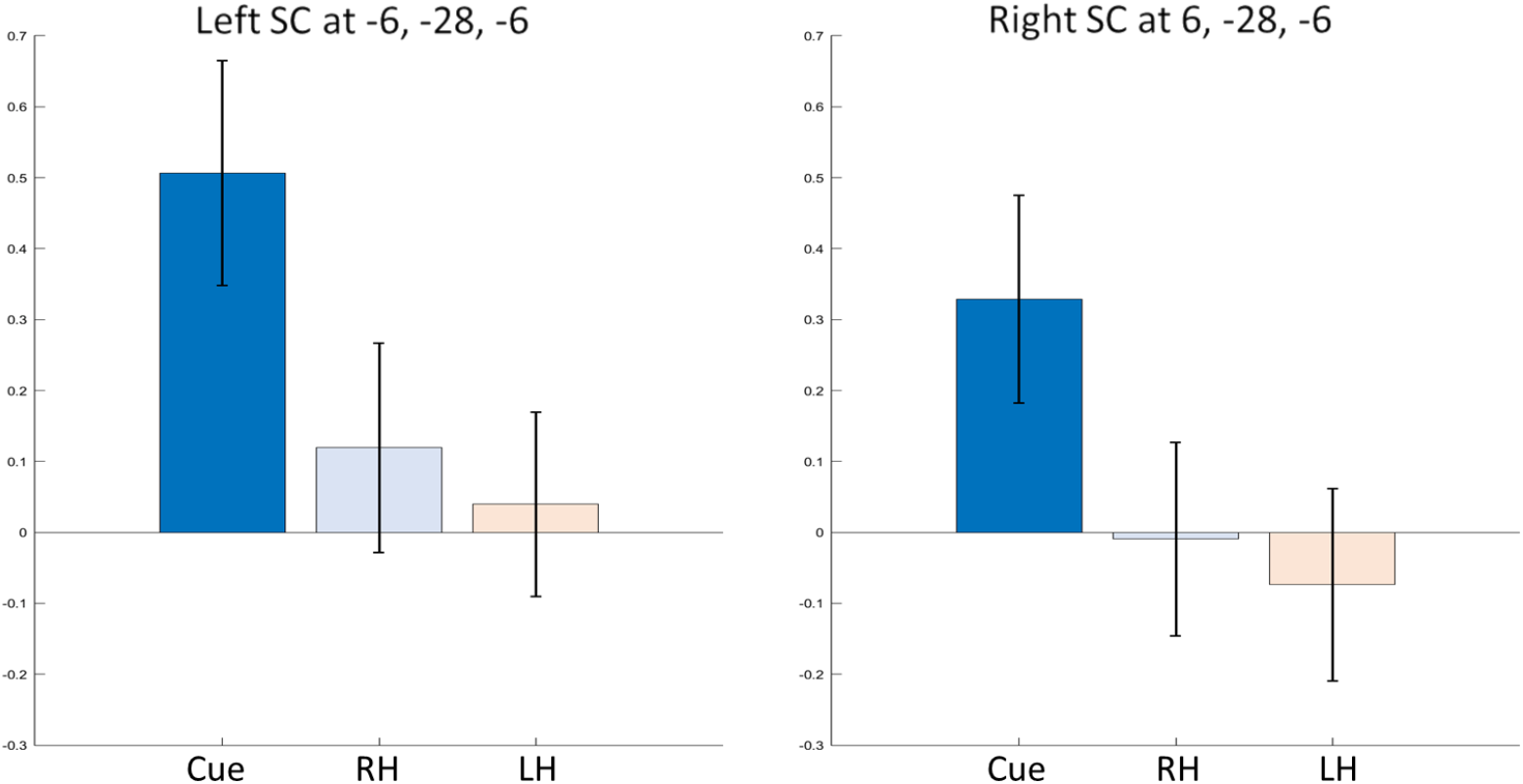
GLM contrast estimates at VOI locations from the HCP motor task dataset. Signal contributions from the three conditions – visual cue, right (RH) and left hand (LH) finger tapping at the VOI -6, -28, -6 and its corresponding location in the right SC: 6, -28, -6. Error bars indicate 90% confidence intervals.

We further explored voxel-wise second-level GLM results for the three conditions of interest in the SC with a significance threshold of *p <* 0.001, uncorrected. We observed a strong response for visual cues but found no response for the right- and left hand finger tapping movements in either the left or right SC (Figure 3). These results confirmed and complemented the findings from the localized contrast estimates reported above.

**Figure 3:**
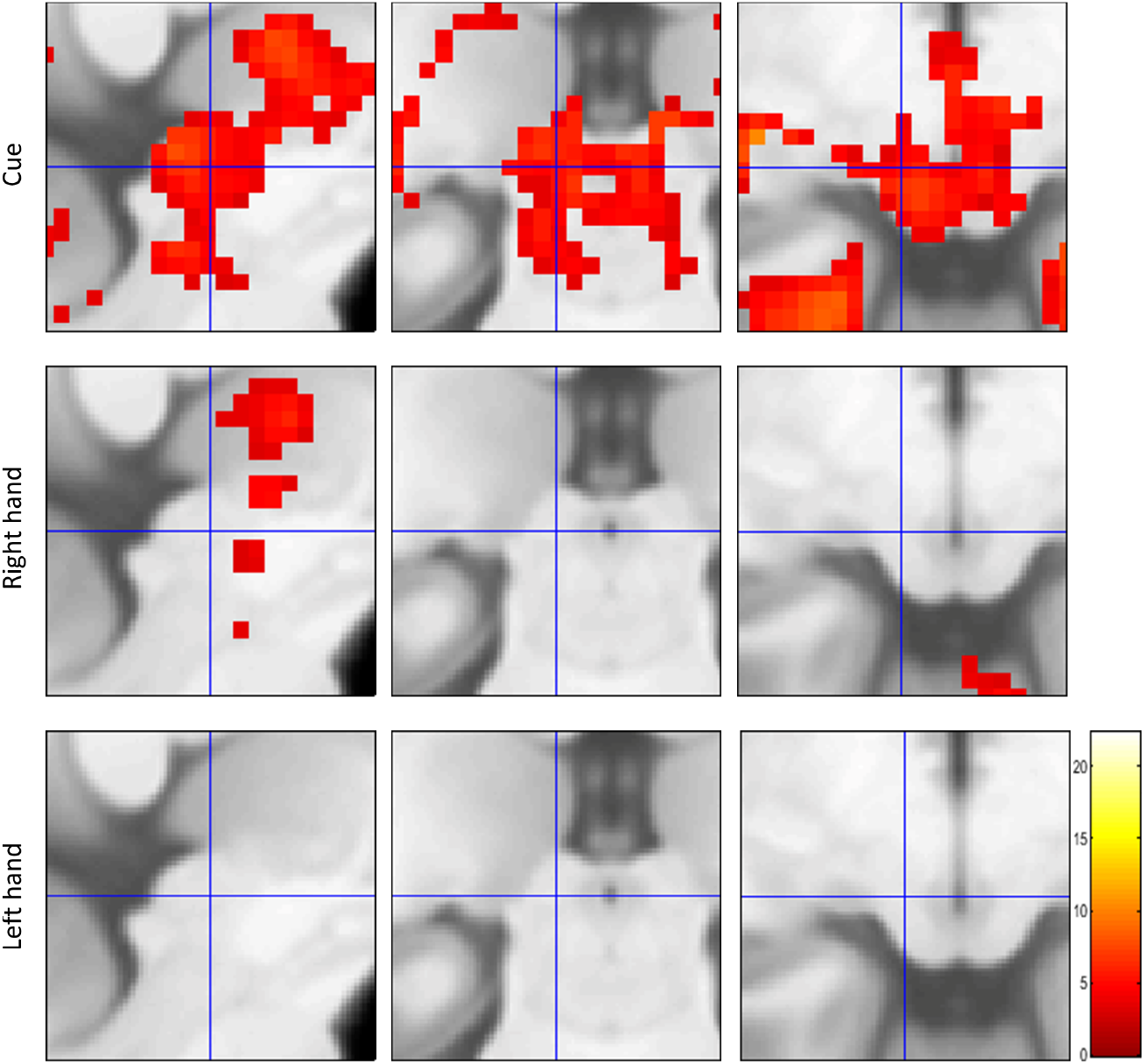
Voxel-wise GLM results in the SC from the HCP motor task dataset. Second-level GLM analysis results from cue, left hand, and right hand finger tapping conditions compared with fixation baseline (*p <* 0.001, uncorrected). The images are centred on the VOI -6, -28, -6. There were no suprathreshold clusters in either left or right SC for the finger tapping conditions. The colourbar shows the range of t-values.

### Simple and complex finger tapping experiment

#### Behavioural data

We further explored our failure to find finger movement signals in the SC in our analysis of the group HCP data with an experiment including simple and complex finger tapping paced by visual and auditory stimuli. The combination of the two experimental factors resulted in four experimental conditions, visual simple finger tapping (VS), auditory simple finger tapping (AS), and visual and auditory complex finger tapping (VC and AC). We measured 4 participants with each participant repeating the experiment on 4 different days. We monitored gaze fixation and finger movements during the fMRI measurements with MRI-compatible cameras. Out of a total of 55 intact and complete fMRI runs from the 4 subjects, eye videos from 44 runs were available for the behavioural analysis, 12, 10, 12, and 10 videos respectively, from the 4 subjects (please see methods section below for details). Each run consisted of 16 experimental condition blocks with 40 button presses per block, resulting in 640 button presses per run. The average number of unwanted saccades per run, breaking gaze fixation, varied between 0 and 10 in subjects 1,2 and 3. In the 4^th^ subject the average number of unwanted saccades varied between 3 and 29. In runs with only simple button presses, saccades per run varied between 0 and 2, whereas in runs with complex button presses, saccades per run varied between 0 and 4, in the first 3 subjects. In the 4^th^ subject, in runs with simple button presses the number of saccades per run varied between 7 and 29 whereas in runs with complex button presses it varied between 24 and 28.

#### Bold fMRI results

To ensure that the experimental paradigm elicited a response in cortical motor areas and thus confirm the validity of our approach we first examined results from the hand knob in M1 (primary motor cortex), PMd (dorsal pre-motor cortex), and PMv (ventral pre-motor cortex). We visually localised the hand knob in M1 and identified the local maxima at this anatomical location. Figure 4 shows the co-ordinates for the local maxima in all four conditions at the hand knob in M1 for each subject thresholded at *p <* 0.05, FWE-corrected. We searched for the local maxima close to the peak signal co-ordinates for PMd and PMv reported in the meta-analysis by Mayka et al. (2006). We then verified that these individual local maxima lay within the boundaries of the respective structures as described in the meta-analysis by Mayka et al. (2006). Local maxima for PMd and PMv across all conditions are reported in table 1, thresholded at *p <* 0.05, FWE-corrected. We found a positive response in all four task conditions in M1, PMd, and PMv in each subject.

**Figure 4:**
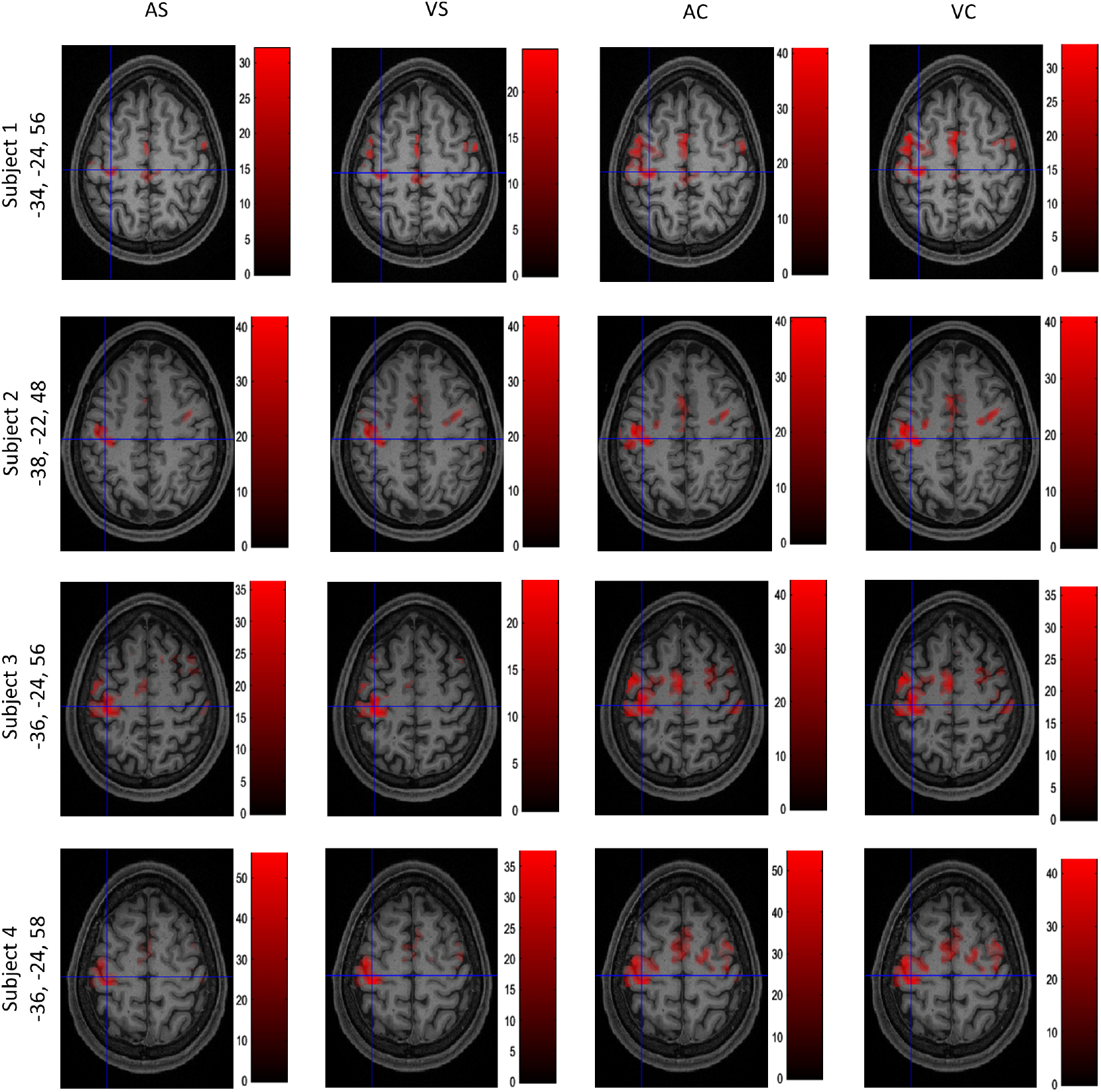
Local maxima in M1 in all subjects and conditions. Activation maps with crosshairs centred on the local maximum in the hand knob area of the left M1 across all four subjects. T-contrasts for the four experimental conditions against baseline: auditory simple (AS), auditory complex (AC), visual simple (VS), and visual complex (VC) thresholded at *p <* 0.05, FWE-corrected. Colourbars show the range of t-values.

**Table 1:** MNI co-ordinates for local maxima in the left PMd and PMv across all four finger tapping conditions, auditory simple, auditory complex, visual simple and visual complex, thresholded at *p <* 0.05, FWE-corrected. The respective range of t-values is reported across the 4 conditions.

| Subject | PMd MNI x, y, z | t-values | PMv MNI x, y, z | t-values |
| --- | --- | --- | --- | --- |
| 1 | -38, -18, 48 | 10.49 - 19.53 | -58, 0, 32 | 14.39 - 27.12 |
| 2 | -36, -16, 44 | 22.26 - 32.12 | -56, 6, 28 | 7.14 - 16.92 |
| 3 | -36, -18, 58 | 21.39 - 42.46 | -58, 2, 22 | 10.22 - 24.48 |
| 4 | -38, -18, 50 | 37.58 - 49.35 | -54, 6, 22 | 19.22 - 32.80 |

In the current experiment we added a complex finger tapping paradigm that was not part of the HCP dataset, because we expected a differential response in comparison to simple finger tapping. In agreement with our expectations and previous reports, we found a significant difference in M1, PMd and PMv (Figure 5). The local maxima reported in the figure were verified to lie within the boundaries of M1, PMd, and PMv as reported in the meta analysis by Mayka et al. (2006).

**Figure 5:**
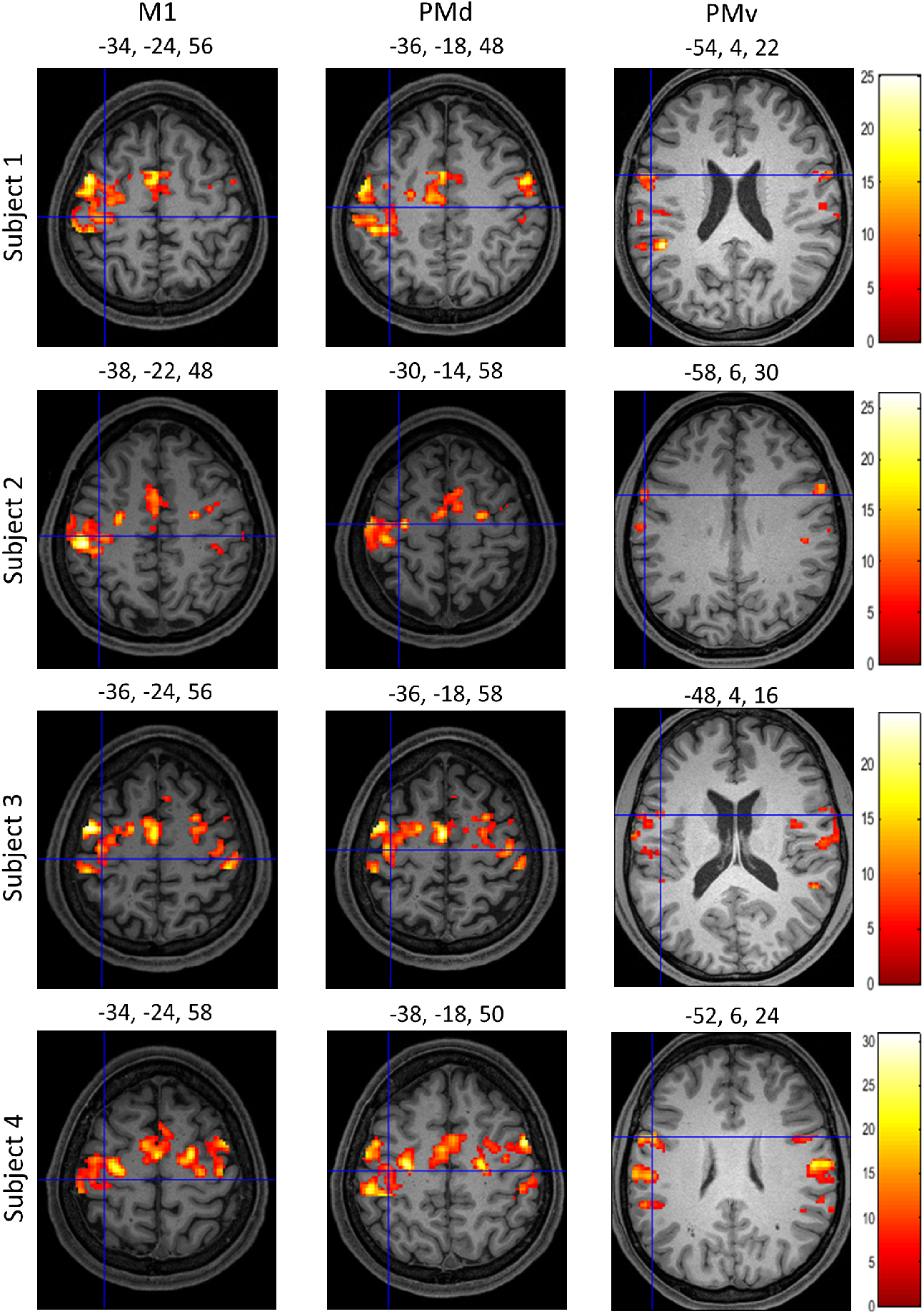
Signal increases in complex compared to simple finger tapping. Activation maps for a differential T-contrast with crosshairs centred on the local maxima in the left hand knob area of M1, PMd, and PMv in the 4 subjects (*p <* 0.05, FWE-corrected). Colourbars show the range of t-values.

We then analysed GLM contrast estimates for each subject at the VOI -6, -28, -6 and in the corresponding location in the right SC for all four experimental conditions and visual cues (Figure 6). Visual cues were associated with a significant positive signal increase in all four subjects, while none of the four finger tapping conditions showed a positive signal in any subject.

**Figure 6:**
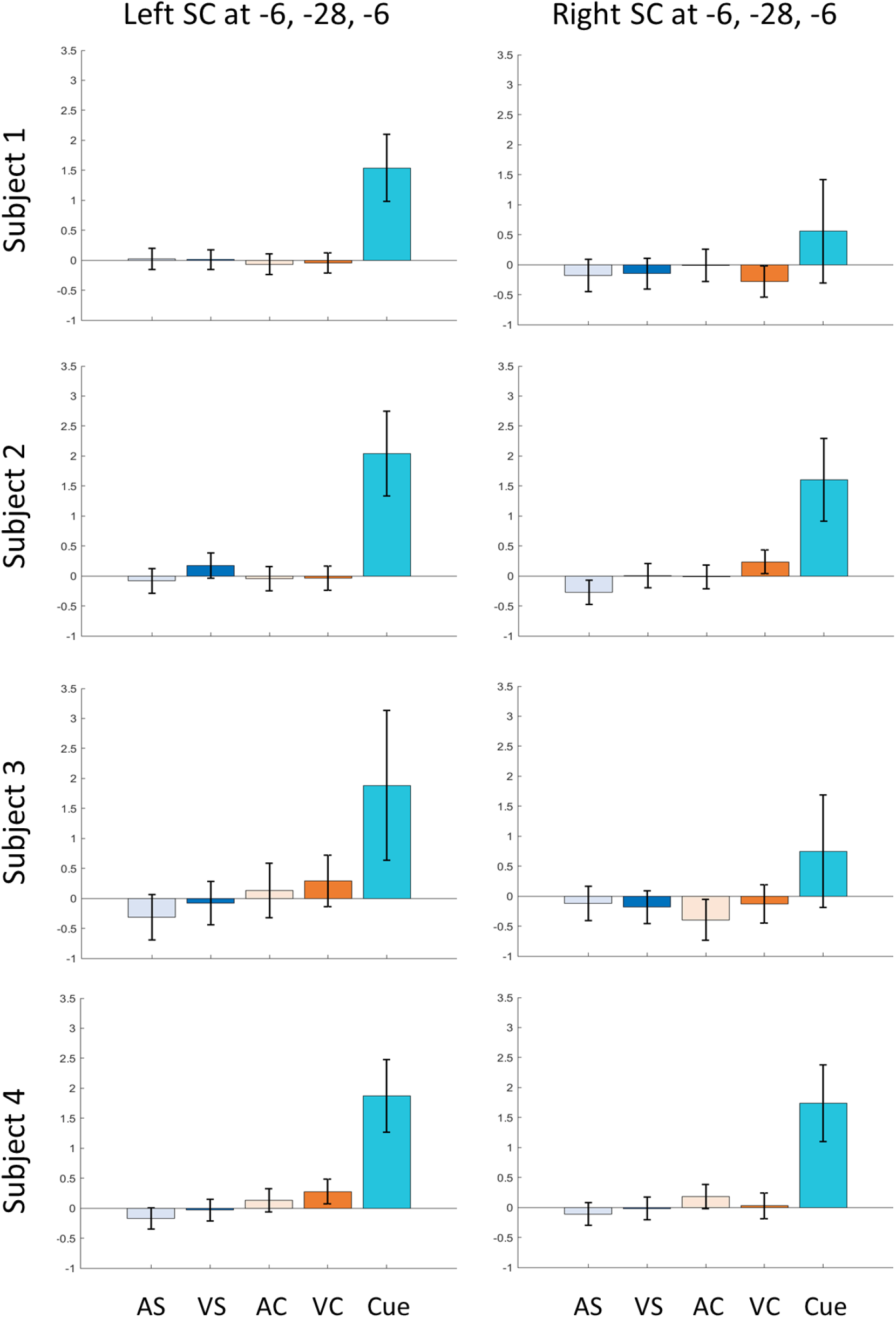
Contrast estimates at the a priori VOI in SC. GLM contrast estimates for the four tapping conditions and visual cues against baseline for each subject. AS: auditory simple, VS: visual simple, AC: auditory complex, VC: visual complex. The error bars indicate 90% confidence intervals.

We further explored voxel-wise GLM results for visual cues and the four conditions of interest in the SC with a significance threshold of *p <* 0.001, uncorrected. A t-contrast against baseline for the cue condition revealed significant clusters in 3 out of 4 subjects (Figure 7). We found no response in the SC above the threshold for any finger tapping condition in t-contrasts against baseline. We further explored the finger tapping conditions using an F-contrast that would have revealed positive or negative signals in any of the four finger tapping conditions, again without positive results.

**Figure 7:**
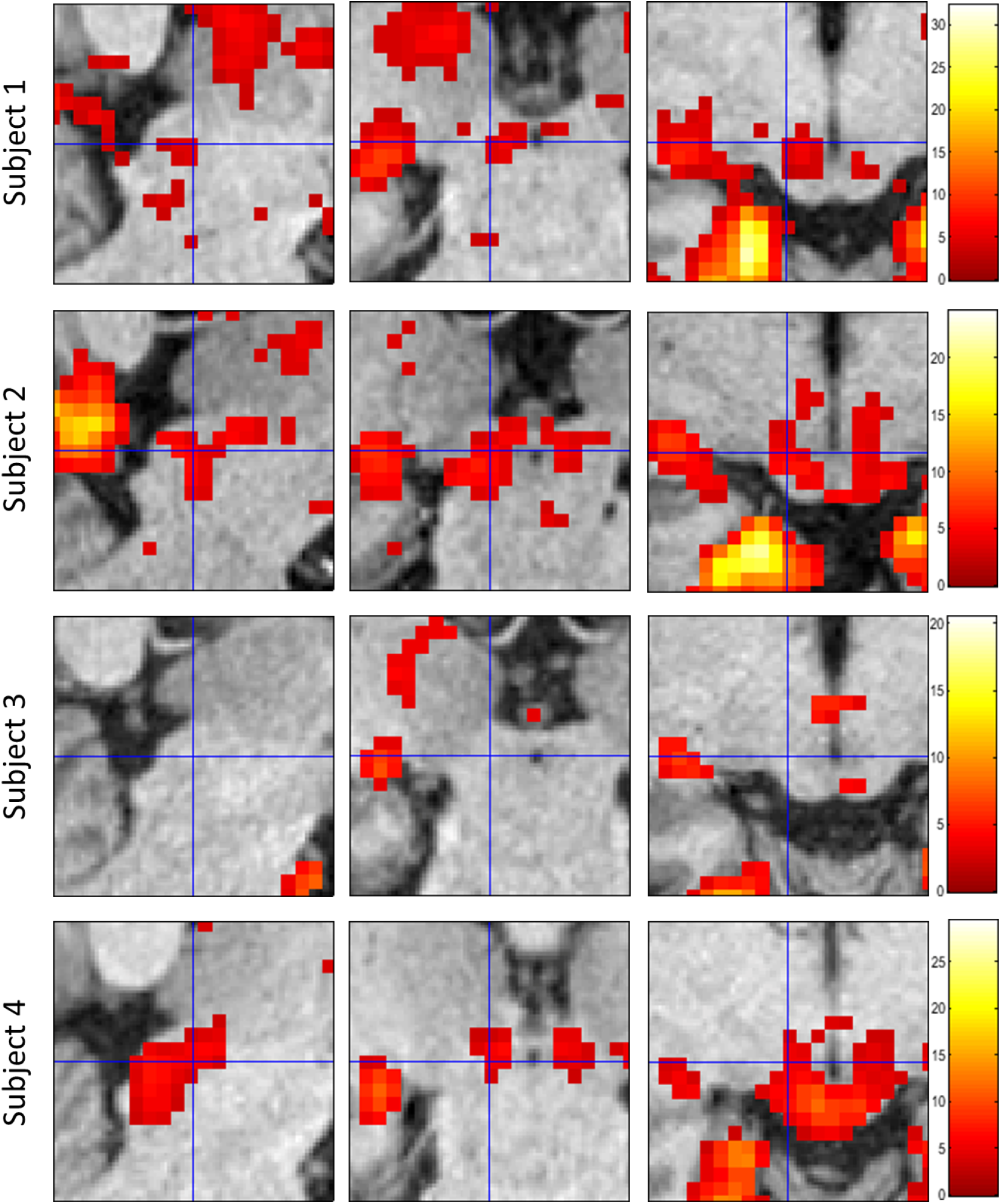
Voxel-wise GLM results for cues vs baseline. Suprathreshold voxels are shown in sagittal, coronal, and transverse sections of the SC centred on the VOI -6, -28, -6 at a threshold of *p <* 0.001, uncorrected. Colourbars show the range of t-values.

We then examined signals in the four conditions in the SC, PMd, and PMv using detailed time courses (Figure 8). The within-subject mean time courses in the SC have been extracted for the a priori VOI for each subject whereas timecourses from PMd and PMv have been extracted for each subject’s individual PMd and PMv voxels as reported above (Table 1). PMd and PMv showed clear signal increases in good correspondence with block onset and duration (Figure 8). We found the highest peaks above baseline in the PMd. We found higher peaks for complex tapping tasks compared to simple tapping in all 4 subjects in PMd and PMv. In stark contrast to time courses in PMd and PMv, there was no indication of any signal increase above baseline in the left or right SC.

**Figure 8:**
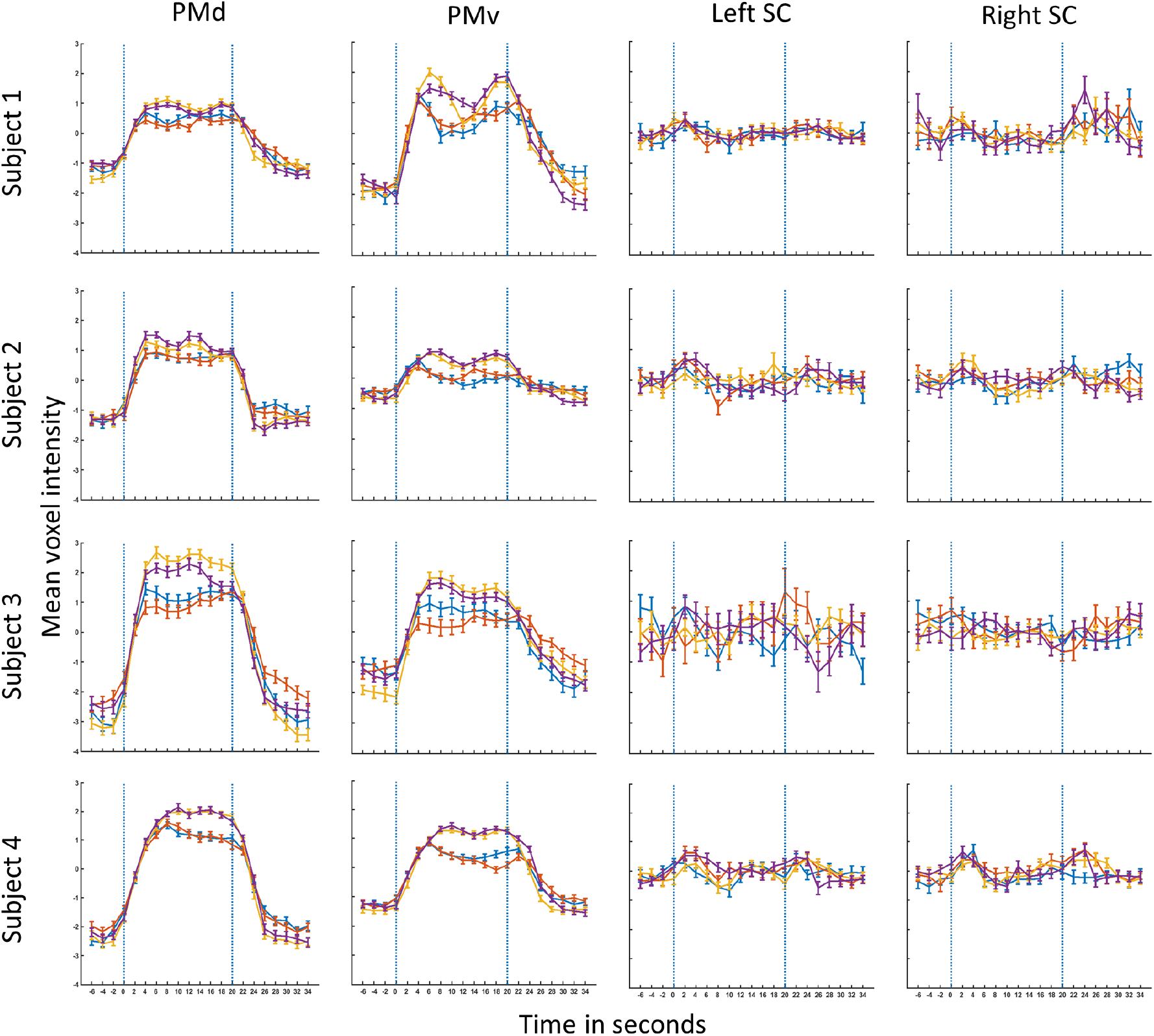
Signal time courses for each subject and condition in PMd, PMv, and SC. PMd and PMv time courses have been extracted for the voxels reported in table 1. SC time courses have been extracted for the VOI (−6, -28, -6) and its corresponding location in the right SC (6, -28, -6). The signal time courses have been extracted for a time window from -6 s relative to block onset to 15 s after the end of the block. The two vertical blue dotted lines indicate the start and end of the block, 0 and 20 s respectively. The error bars indicate standard error (SE). Blue – Auditory simple (AS), Red – Visual Simple (VS), Yellow – Auditory Complex (AC) and Purple – Visual Complex (VC).

## Discussion

We did not find any signals corresponding to finger tapping in the human SC in our analyses. The human SC limb motor region seems to be involved only in reaching guided by visual and tactile stimulation (Linzenbold and Himmelbach, 2012; Himmelbach et al., 2013; Prabhu and Himmelbach, 2022). Given that Isa et al. (2021) and Cooper and McPeek (2021) implicated the SC in various motor tasks and activities across species, it is intriguing that the functional role of the human SC motor region does not include distal finger tapping movements.

The premotor cortices, PMd and PMv, have been implicated in the integration of sensory information towards realisation of sequential movements (Jäncke, Himmelbach et al., 2000; Jäncke, Loose et al., 2000; Horenstein et al., 2009; Ruspantini et al., 2011; Genon et al., 2017; see Witt et al., 2008 for a meta-analysis). Studies by Fries (1985), Borra et al. (2014), and Distler and Hoffmann (2015) described a connection between the premotor cortices and the SC limb motor region. The anatomical connection between the SC and cortical areas that are involved in the execution of sequential finger movements, prompted us to investigate if the SC also plays a role in such movements. We first analysed a large, publicly available dataset where we found activation clusters in M1, PMd, and PMv corresponding to finger tapping. There was no evidence for finger tapping related activity in the SC, despite of strong responses to visual stimulation in the same measurements. We then performed an experiment in 4 subjects with each subject repeating the experiment 4 times where we saw activation in M1, PMv, and PMd corresponding to simple and complex finger tapping sequences and even differentiating between simple and complex tapping. Again, we found no signals in the SC corresponding to either simple or complex finger tapping sequences, although visual cues elicited a response similar to our analysis of the HCP dataset. We replicated this pattern of results in all 4 participants with a large number of repetitions across different sessions and days.

The results of our preceding study (Prabhu and Himmelbach, 2022) clearly showed a response in the SC for individual button presses, i.e. finger tapping with a button. In contrast, our current study did not provide any evidence for a response in the SC. In our earlier study, finger tapping movements were discrete and infrequent and instructed by an odd-one-out event. In our current study, finger movements were repetitive and not individually instructed. The simple visual and auditory pacing stimuli in our current experimental measurements were fundamentally different from the odd-one-out stimuli in our previous experiment (Prabhu and Himmelbach, 2022) with respect to their information value. Results from a study using electrical microstimulation in macaques might be of relevance for the results of the current study. Philipp and Hoffmann (2014) induced upper limb twitches, lifting, and extending movements by electrical stimulation of the deep, posterior-lateral SC, i.e. the SC upper limb motor region. Interestingly, the type of induced movements in the three animals depended on prior experience and training. The animal with extensive prior training in reaching paradigms showed the highest proportion of actual reaching movements upon electrical stimulation (Philipp and Hoffmann, 2014). It seems like the SC does not directly initiate reaching or tapping but is embedded in a larger network that gates and modulates the motor functions of the SC. The SC can be seen as a fast-responding system with a large variability of the behavioural output across species (Cooper and McPeek, 2021). In comparison to orientation behaviour, evasive movements, or reaching, sequential finger-tapping is a phylogenetically young behaviour without an immediate goal related to environmental demands. Further research on motor functions of the SC might therefore investigate finger movements that are embedded in a behavioural context with more ecological relevance, e.g. grasping and catching.

While positive findings in fMRI studies are frequently confronted with suspicions of analytic flexibility (Carp, 2012, 2012a), negative findings in fMRI raise concerns on the sensitivity and detection power of the respective paradigms and analyses. We are convinced that our negative findings cannot be explained by an insufficient experimental design or analysis. For the HCP data analysis as well as for our own experimental measurements, activation clusters in M1, PMd, and PMv showed that the paradigms and analyses were adequate to reveal responses in motor areas corresponding to finger tapping. Activation clusters for the visual cue conditions from both analyses indicated that the paradigms and sequences were suited for signal detection at the SC. With the HCP analysis, we showed that the SC does not play a role in the execution of simple finger tapping movements. With the second study, we replicated negative findings from the analysis of the HCP dataset for simple finger tapping in four individual subjects. In addition, we found that this result does not change with more complex but overlearned finger tapping sequences. The inspection of our descriptive data clearly showed an overlap of motor related signals at the SC with baseline signal levels in each analysis. While non-significant, inferential statistics cannot provide conclusive evidence for a null finding, the descriptive data from both analyses convinced us of a true null finding at the human SC motor region.

## Methods

### Human Connectome Project data

#### Subject details

We analysed 130 subjects from the HCP database that had a complete dataset for the motor task (73 females and 57 males, age range 22 - 36 years, 10 left-handed and 120 right-handed) with normal or corrected-to-normal visual acuity.

#### Experimental setup and paradigm

The HCP motor task consisted of 13 blocks per run, 10 movement blocks and 3 interspersed fixation blocks. Each movement block lasted 12 s and consisted of 10 movements. A movement block could consist of tapping the fingers of the left hand or the right hand, or squeezing of the left or right toe, or a movement of the tongue depending on a visual cue that was shown for 3 seconds just before the start of the block. The 10 movement blocks were split into 2 tongue movement blocks, 4 hand movement blocks (2 right and 2 left) and 4 foot movement blocks (2 right and 2 left). The 3 fixation blocks in a run lasted 15s in duration. 2 runs with 13 blocks each were measured from each subject (Barch et al., 2013).

#### fMRI data acquisition

Data acquisition was performed at Washington University using a modified 3T Siemens Skyra MRI scanner. A 32-channel head coil was used for the whole-brain acquisition of functional measurements (Gradient echo - EPI) with a TR = 720 ms, TE = 33.1 ms, flip angle = 52^*◦*^, FOV = 208, 72 slices, with a voxel resolution of 2.0 mm isotropic, and a multi-band acceleration factor of 8. The two runs for the motor tasks were measured with opposing phase-encoding directions of right to left and left to right respectively. The structural images were acquired with a 3D MPRAGE sequence TR = 2400s, TE = 2.22ms, flip angle = 8^*◦*^, FOV = 224 × 224 mm (320 × 320 matrix) and a voxel resolution of 0.7 mm isotropic. For further details on imaging protocols of the HCP, please refer to https://www.humanconnectome.org/hcp-protocols-ya-3t-imaging

#### A priori definition of the volume of interest

In a preceding study on reaching-related activity in the SC, we observed BOLD activity with finger tapping for button presses in response to oddball stimuli that were shown in control conditions (Prabhu and Himmelbach, 2022). We extracted data from a 3×3×3 grid of 27 voxels in the posterior-lateral SC limb motor region in the left and right SC. We then identified the voxel with the highest t-value in the finger tapping/button press condition and averaged these voxel co-ordinates across subjects. The resultant voxel was located at -6, -28, -6 (MNI co-ordinates; left SC). We used this voxel as the volume of interest for the analyses in the current study. In addition, we also examined results at the corresponding location in the right SC at 6, -28, -6.

#### Data pre-processing

We used the minimally pre-processed dataset that was available with the HCP consortium. This data had already undergone pre-processing procedures like re-alignment, co-registration and normalisation except for smoothing (for details refer to Glasser et al., 2013). We smoothed the data with a full-width half-maximum (FWHM) Gaussian kernel of 3.0 mm.

#### fMRI data analysis

We defined a first-level model with block onsets for the cue, left hand finger movements, right hand finger movements, left toe movements, right toe movements, and tongue movements with block durations of 3 s for the cues and 12 s for the movement conditions. These regressors were convolved with a modified hemodynamic response function with a shorter onset-to-peak time of 4 s that better fits neurovascular response dynamics in the SC (Wall et al., 2009). The HCP data comprises 12 re-alignment parameters out of which the first 6, corresponding to 3 translations and 3 rotations of head movements, were used as regressors of no-interest. Contrast images were then calculated with each condition in the experiment compared against the baseline. The second-level model was built with the contrasts of interest – cue, left hand and right hand finger movements. We analysed group GLM contrast estimates at the VOI mentioned above. We expected positive signal increases and considered results to be significant if the lower bound of the two-sided 90% confidence interval, which amounted to 95% confidence for our uni-directional hypothesis, did not include the zero baseline. Additionally, we conducted a voxel-wise analysis of cortical regions and SC surrounding the VOI. Results were considered to be significant surviving a voxel-level threshold of 0.05 FWE-corrected for cortical results and 0.001 uncorrected for results in the SC. Please note that the more conservative threshold for cortical data in combination with the more lenient threshold for SC data worked against our conclusion of a null-finding at the SC with positive signal detection in cortical motor areas.

### Simple and complex finger tapping experiment

#### Subject details

We conducted the study using 3T BOLD fMRI with four right-handed healthy volunteers (2 females, 2 males, age range 25 - 32 years) with normal or corrected-to-normal visual acuity. Each subject performed the experiment four times with one session, comprising four runs, per day. The experiments were conducted with the approval of the local ethical committee and as per the ethical standards established by the 1964 declaration of Helsinki in its latest version. Informed consent was obtained from all participants.

#### Experimental setup and paradigm

Subjects held a button box in their right hand with four buttons and wore headphones. We placed cushions made especially for fMRI experiments (NoMoCo Pillow, Inc., La Jolla, USA) in the head-coil of the scanner to immobilise the head of the subjects. Two MR-compatible cameras (MRC systems GmbH, Germany) operating at 30 Hz, one to monitor finger movements and the other to monitor eye movements, were attached to a stand that was placed across the waist of the subject. Fibre optic cables carrying light from three LEDs; a white fixation/pacing LED, a green cue LED and a red cue LED were attached to the uppermost central part of the stand. Subjects saw the optic fibre endings through a mirror attached to the head-coil. Auditory pacing stimuli were delivered through the headphones. The entire experimental paradigm including stimulus delivery and control of timing was implemented on a ‘MBED LPC 1768’ microcontroller (Arm Limited). All analyses were carried out using MATLAB (version R2018a, The MathWorks Inc., Natick, MA, USA).

We trained subjects to pace finger tapping in accordance with visual or auditory pacing stimuli, just before the experiment began, until they successfully matched two blocks of stimuli with finger tapping in a row. Our subjects accomplished this in a maximum of 10 minutes. Finger tapping could have been of two kinds – simple or complex finger tapping. Simple finger tapping involved pressing one of the four buttons paced by cues while complex finger tapping involved repetitions of the four buttons; 2 times first button (index finger), 4 times second button (middle finger), 1 time third button (ring finger), 3 times fourth button (little finger) followed by finger tapping in the reverse order starting from the little finger (task adapted from Kuhtz-Buschbeck et al., 2003). The simple and complex finger tapping sequence blocks were informed by a cue of 1 s duration. A green LED cued the simple finger tapping block and a red LED cued the complex finger tapping block. For the simple finger tapping task, the green LED also cued the finger (any of the four) to be used in the impending block with the number of times it blinked i.e. one blink for the first finger (index finger), two blinks for the second finger (middle finger) and so on. Block pacing stimuli were of two kinds - visual or auditory, and were presented at 2 Hz with either a blinking white LED or a ‘beep’ delivered through headphones, respectively. The scanner room was completely darkened by covering windows and scanner displays with black opaque film. We set the brightness of the fixation light to a level just enough to discern it from the surroundings hence ensuring that it did not illuminate the workspace. The subjects looked at the central fixation light throughout the experimental run time of 8 minutes and 50 seconds per run. The two modalities, visual and auditory, along with the two types of responses, simple and complex finger tapping, were combined in a 2×2 factorial design. This resulted in 4 conditions: visual simple (VS), visual complex (VC), auditory simple (AS), and auditory complex (AC), two of them being part of each run, with one common factor. The four resulting runs AS-VS, AC-VC, AS-AC, and VS-VC were ordered in a Latin-square arrangement across the 16 runs within each subject, to control for sequence effects. The two conditions in each run alternated as 20 s blocks of 40 trials, each experimental block interleaved by a fixation baseline of 15.5 s.

#### Analysis of saccade data

Eye videos were converted into eye traces with DeepLabCut (Mathis et al., 2018). Saccade data with a threshold of 2° visual angle were extracted from these eye traces. Synchronisation errors in the video frame grabber meant that several videos were unreliable for analysis. Missing frames in the videos did not allow for a temporal analysis of saccades in the video. Hence, we analysed and reported the total number of saccades per subject per run.

#### fMRI data acquisition

We performed the measurements using a 3T Siemens Prisma scanner with a 64-channel head-coil. 258 functional images (T2* weighted) were acquired as part of each run – interleaved BOLD imaging of 28 slices in a coronal orientation covering the pre-motor areas and the brainstem, bilaterally. The images were acquired at a resolution of 2 mm isotropic with a TR = 2360 ms, TE = 35.0 ms, flip angle = 90^*◦*^, FOV = 180 mm x 180 mm (90 × 90 matrix) and a phase encoding direction of foot to head. At the end of each imaging session a whole-brain EPI image was acquired to aid in co-registration with the functional partial volumes. A structural image (T1 weighted) was acquired per subject using an MP-RAGE sequence with a resolution of 0.8mm isotropic and TR = 2400 ms, TE = 2.22 ms, flip angle: 8^*◦*^, FOV = 240 mm x 256 mm (300 × 320 matrix).

#### Pre-processing of MRI data

Pre-processing and the following GLM analysis were carried out using Statistical Parametric Mapping (SPM12, Wellcome Trust Centre for Neuroimaging, London, UK), which was implemented in MATLAB (version R2018a, The MathWorks Inc., Natick, MA, USA). During an inspection of the images converted into the Nifti format we observed that some runs had distorted images. Such runs with image artifacts, along with those where button press timings were not reliable due to technical errors, were discarded from further analysis. The number of runs that underwent further analysis were all 16 runs from the first subject, 12 runs from the second, 12 runs from the third, and 15 runs from the last subject (55 runs in total). We deleted the first five images from all runs to allow scanner signals to reach a steady state. The rest of the images underwent a re-alignment procedure as defined in SPM, correcting for motion. The co-registration procedure was performed as follows. First, the mean EPI scans from each of the sessions in a subject were co-registered with the whole-brain EPI scan of the respective session. Next, the whole-brain EPI scans from all sessions of a subject were co-registered with the whole-brain EPI image from the first session of the subject. Lastly, the whole-brain EPI scan from the first session was co-registered with the structural image, followed by an inspection for conformity between the mean functional scans across sessions with the respective structural image. Mis-aligned images were first manually oriented with the structural images before undergoing normalisation with a MNI152 (Montreal Neurological Institute) template. This resulted in good co-registration between the functional images, structural images, and the template. The images were then smoothed with a full-width half-maximum (FWHM) Gaussian kernel of 3mm.

#### fMRI data analysis

A first-level analysis for each subject was performed using SPM12 with a dataset that included all sessions from different days. The first six motion parameters from the re-alignment step of the pre-processing stage were used as regressors of no-interest. The first level model was designed with block onsets from each condition: Visual simple (VS), Visual complex (VC), Auditory simple (AS) and Auditory complex (AC) along with block durations, and onset and duration of visual cues as regressors of interest. These regressors were convolved with a modified hemodynamic response function with a shorter onset-to-peak time of 4s that better fits neurovascular response dynamics in the SC than the canonical response function implemented in SPM12 (Wall et al., 2009). Contrasts were built to examine results for each of the conditions against the baseline, followed by a contrast of simple tasks against the corresponding complex conditions. An Omnibus F-contrast was also built to examine activity in the SC from all conditions together. We analysed within-subject GLM contrast estimates at the VOI mentioned above. We expected positive signal increases and considered results to be significant if the lower bound of the two-sided 90% confidence interval, which amounted to 95% confidence for our uni-directional hypothesis, did not include the zero baseline. Additionally, we conducted a voxel-wise analysis of cortical regions and SC surrounding the VOI. Results were considered to be significant surviving a voxel-level threshold of 0.05 FWE-corrected for cortical results and 0.001 uncorrected for results in the SC.

To examine voxel signal time courses we extracted raw time courses spanning all sessions from a subject. The raw time courses were then de-meaned across sessions, regressed with motion parameters and high-pass filtered at 128 Hz. The time courses were interpolated to 0.1 s and later re-sampled to 2 s intervals aligned to the start of each experimental block. The time courses were then separated by conditions using their respective block onsets. A within-subject mean time course with standard errors was computed for each condition.

## Notes

### Competing Interest Statement

The authors have declared no competing interest.

